# Rare Copy Number Variant analysis in case-control studies using SNP Array Data: a scalable and automated data analysis pipeline

**DOI:** 10.1101/2024.03.13.584428

**Authors:** Haydee Artaza, Ksenia Lavrichenko, Anette S.B. Wolff, Ellen C. Røyrvik, Marc Vaudel, Stefan Johansson

**Affiliations:** Department of Clinical Science, University of Bergen, Bergen, Bergen, Norway; K.G. Jebsen Center for Autoimmune Diseases, University of Bergen, Bergen, Norway; Department of Medical Genetics, Oslo University Hospital, Oslo, Norway; Department of Medicine, Haukeland University Hospital, Bergen, Norway; Department of Genetics and Bioinformatics, Norwegian Institute of Public Health, Bergen, Norway; Mohn Center for Diabetes Precision Medicine, Department of Clinical Science, University of Bergen, Bergen, Norway; Department of Medical Genetics, Haukeland University Hospital, Bergen, Norway

**Keywords:** Copy number variant (CNV), calls detection, quality control, burden analysis, enrichment analysis, rare variants analysis, snakemake

## Abstract

**Background:** Rare copy number variants (CNVs) significantly influence the human genome and may contribute to disease susceptibility. High-throughput SNP genotyping platforms provide data that can be used for CNV detection, but it requires the complex pipelining of bioinformatic tools. Here, we propose a flexible bioinformatic pipeline for rare CNV analysis from human SNP array data.

**Results:** The pipeline performs two major tasks: (1) CNV detection and quality control, and (2) rare CNV analysis. It is implemented in Snakemake following a rule-based structure that enables automation and scalability while maintaining flexibility.

**Conclusions:** Our pipeline automates the detection and analysis of rare CNVs. It implements a rigorous CNV quality control, assesses the frequencies of these rare CNVs in patients versus controls, and evaluates the impact of CNVs on specific genes or pathways. We hence aim to provide an efficient yet flexible bioinformatic framework to investigate rare CNVs in biomedical research.

## Background

Copy number variation (CNV), defined here as deletions and duplications of chromosomal segments larger than 1 kb, are a major source of genetic variation between individuals and are an essential factor in many complex diseases, including mental illness, developmental disorders, and cancer[1]. In particular, distinct large (> 1000 kb) CNVs have been linked to rare disease phenotypes, and they may contribute to common polygenic diseases[2]. High-throughput single nucleotide polymorphism (SNP)-array technologies allow the investigation of CNVs at a genome-wide scale[3].

A large number of studies have investigated rare CNVs using micro-array based genotyping data[4–10]. These investigations typically involve intricate procedures, necessitating multiple analyses, careful choice of software, calibration of sensitivity to parameters and their thresholds, and execution setting. Computational and scientific outcomes therefore hinge upon automation and thorough documentation of implementation specifics. Standardized basic protocols for calling CNV and performing association tests have been proposed by others, such as in Lin *et al*.[11], however a comprehensive simple-to-use bioinformatic implementation has not been provided.

Conducting a case-control study based on rare CNVs involves several critical steps: (1) CNV detection, (2) quality control, (3) burden analysis, and (4) gene-set enrichment analysis. High-throughput next-generation genotyping technologies, commonly employed in genome-wide association studies (GWAS), provide the signal intensity data necessary for CNV detection. Subsequently, tools like PennCNV[12] and Plink[13] are typically used for the case-control analysis of CNV, focusing on individual-based CNV calls, and rare CNVs, respectively. Conducting such analyses therefore requires adeptly applying and coordinating multiple advanced bioinformatic softwares, but to the best of our knowledge a bioinformatic pipeline implementing rare CNV analysis in a structured, flexible, and scalable manner remains missing.

In this work, we present a generic bioinformatic solution for identifying rare CNVs in case-control studies. Our main goal is to provide a flexible tool that enables users to conduct rare CNV analysis using SNP array data from different case-control studies.

## Implementation

We have employed the Snakemake workflow[14] engine to construct a robust pipeline for rare CNV analysis. The code is modular and rule-based, leveraging Snakemake’s module statement (Figure 1). Notably, if input files are missing or an execution error occurs, the pipeline automatically deletes any generated output files to maintain consistency. The rule-based structure enables automation while maintaining flexibility: the pipeline can be modified according to the nature of the study through parameters, software, or the addition of custom code. The pipeline further provides configuration files and execution logs, along with diagnostic plots produced using the R programming language[15]. Dependencies are managed using Conda[16]. The pipeline is open source, released as a permissive MIT license[17], and the code is available along with documentation from our RareCNVsAnalysis GitHub repository[18]. Additionally the working Docker version of the pipeline is available (currently as a branch in the repository), with ongoing work to test and refine the relevant parts of documentation. Detailed information regarding configuration files, input and output formats and contents for each module and rules, are described in the pipeline guide, available for download from RareCNVsAnalysis Github repository in manual/Rare_CNVs_pipeline_guide.pdf.

**Figure 1.**
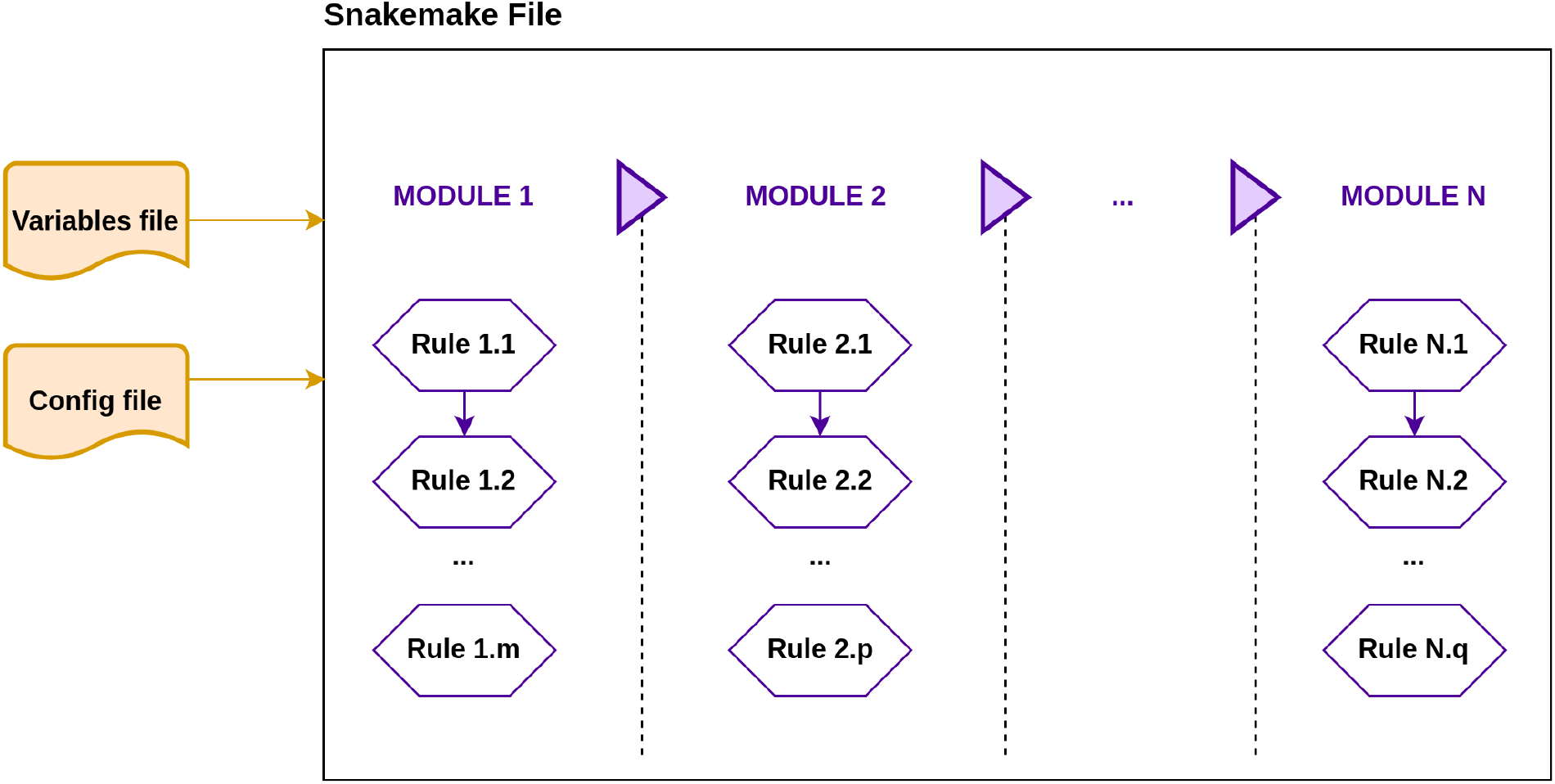
Pipeline structure based on snakemake modules. Our pipeline is organized in modules, each module contains one or more rules. Modules and rules can be modified, added or removed according to the analysis requirements. The list of modules should be included in the snakemake executable file and the description of variables, files and paths should be included in the variables and config files.

### CNV calling and quality control analysis

The initial step of the pipeline performs CNV detection and quality control (Figure 2). It takes the SNP-array genotyping data as input, containing the signal intensity values (Log R Ratio and B Allele Frequency) for all markers in all samples in text format (Pipeline guide, Module Data Conversion). The PennCNV tool offers a user guide to prepare signal intensity files from different array technologies such as Illumina and Affymetrix[19] in the Overview of Input File Formats section. Pipeline processes the signal intensity values extracted from Illumina reports. However, this rule can be readily customized to accommodate alternative signal intensity formats. The cohort-wide signal intensity file is subsequently processed to generate an individual signal intensity file per sample which is utilized in the PennCNV calling process. Additionally, the population frequency of B allele (PFB) and the GCModel files are generated in this step since PennCNV relies on these for accurate CNV detection (more details in https://penncnv.openbioinformatics.org). After CNV detection is carried out, low quality samples are excluded, based on standard genotyping quality metrics: LRR (Log R Ratio), BAF_drift (B Allele Frequency drift), WF (Waviness Factor), and NumCNVs (number of called CNVs). The threshold values for exclusion criteria and the inclusion of other parameters (LRR_mean, LRR_median, LRR_SD, BAF_mean, BAF_median, BAF_SD) can be customized in the pipeline parameters file variables.py (Pipeline Guide, Pipeline Description). Calls detected in challenging genomic regions such as the Human leukocyte antigen (HLA), and the regions near the centromere and the telomeres are considered spurious and are removed[20]. The genomic coordinates of these regions must be contained in external files which will be set into the pipeline configuration file config.json. Finally, the pipeline merges adjacent CNV calls, which often indicate one large CNV (> 500 kb) (Supplementary Figure 1). This analysis yields a set of high quality CNVs which will be the input for further analysis as the rare CNVs.

**Figure 2.**
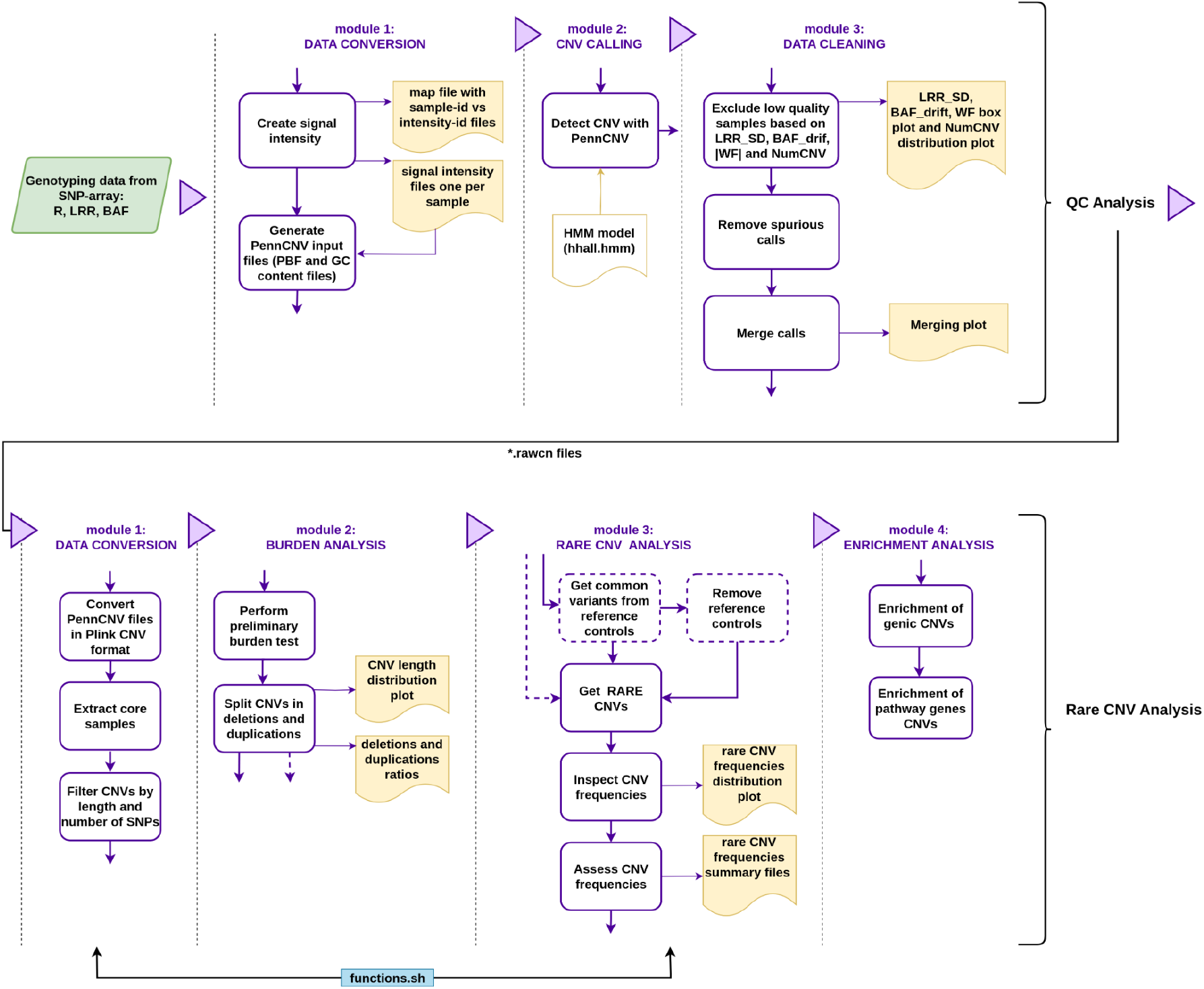
Rare CNVs workflow. The pipeline consists of two major tasks: (1) quality control analysis, which uses the SNP-array genotyping data (green box) as an input to obtain good-quality samples and high-quality calls. (2) rare CNV analysis, which takes samples and calls from the QC pipeline output, and after the data format conversion, performs the burden, rare CNV and enrichment analysis. Black dotted lines split each analysis in their corresponding modules, purple boxes represent a specific task in each module, yellow boxes show representative outputs (files and/or plots), yellow line box represents an external dependency, and the blue box represents external functions used by some modules. Dotted purple boxes are optional tasks which could be easily removed or changed to adapt the pipeline with the study requirements.

### Rare CNVs analysis

The subsequent stage of the pipeline is the analysis of rare CNVs using the calls obtained in the initial step (Figure 2 and Pipeline guide, Rare CNV pipeline). Samples and calls in PennCNV format are first converted to Plink files. Only *core samples*, defined as unrelated and genetically unstratified, are retained, in order to avoid confounding effects[21, 22]. This task requires the users to provide a list with the identifiers of the core samples. These samples can be clustered with a principal component analysis (PCA) or multidimensional scaling (MDS), while the genetic relatedness of the individuals can be based on identity by descent (IBD) analysis^22^ 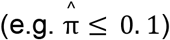. Small CNV calls are usually not reliably detectable by SNP arrays[23, 24], therefore only CNVs larger than 50 kb and covered by more than 5 probes are retained at this stage. Default values can be modified in the parameters file (Pipeline Guide, Pipeline Description).

After sample filtering and CNV size exclusion, a global burden test in cases versus controls is conducted using Plink software. The burden test is performed for four key metrics: (1) number of segments (RATE), (2) proportion of samples with one or more segments (PROP), (3) total kb length spanned (TOTKB), and (4) average segment size (AVGKB). Subsequently, CNVs are divided into deletions and duplications and pooled by length to calculate the CNV frequency in cases versus controls and the CNV distribution within specific length intervals (Supplementary Figure 2). By default, the pipeline defines CNV size thresholds intervals as 50 kb, 100 kb, 200 kb, 500 kb, and 1,000 kb. Users can customize these thresholds in the parameters file (Pipeline Guide, Pipeline Description).

Following the rare CNVs analysis, the pipeline proceeds to extract rare deletions and duplications. This involves identifying common CNVs with frequencies greater than or equal to a user-defined threshold from a subset of healthy control individuals in the study cohort. To calculate the CNV frequency, the Plink overlapping strategy is used. It assigns a specific count to each CNV that represents the number of CNVs (including itself) that overlap with at least 50% of its region. The CNV overlap definition is based on a union intersection approach (Supplementary Figure 3). The subset of healthy individuals involved in the common CNVs identification, are subsequently excluded from further analysis. Using the common CNVs as reference, common variants are filtered out from both the cases and remaining control samples. The next step is to obtain rare CNVs by removing all CNVs with at least 50% overlap with common CNVs. This task is carried out using the BedTools suite[25]. Frequency histograms are generated for quality control of the procedure (Supplementary Figure 4 and 5). It is important to note that our suggested approach to identify rare CNVs can be adapted or modified according to the study strategy. Following this, differences in the frequencies among cases and controls are first assessed for all deletions and duplications, and then, the differences are evaluated for intervals of binned CNV sizes. Summary statistics are generated containing the frequencies for common and rare CNVs in different interval sizes, along with two proportion test statistics and odds ratios (OR) estimation using R version 3.6.3[15], specifically the *stats* and *fmsb* packages. These results are represented graphically as forest plots, with the confidence intervals of frequencies within each CNVs interval size, alongside the associated p-value (Supplementary Figure 6 and 7).

**Figure 3.**
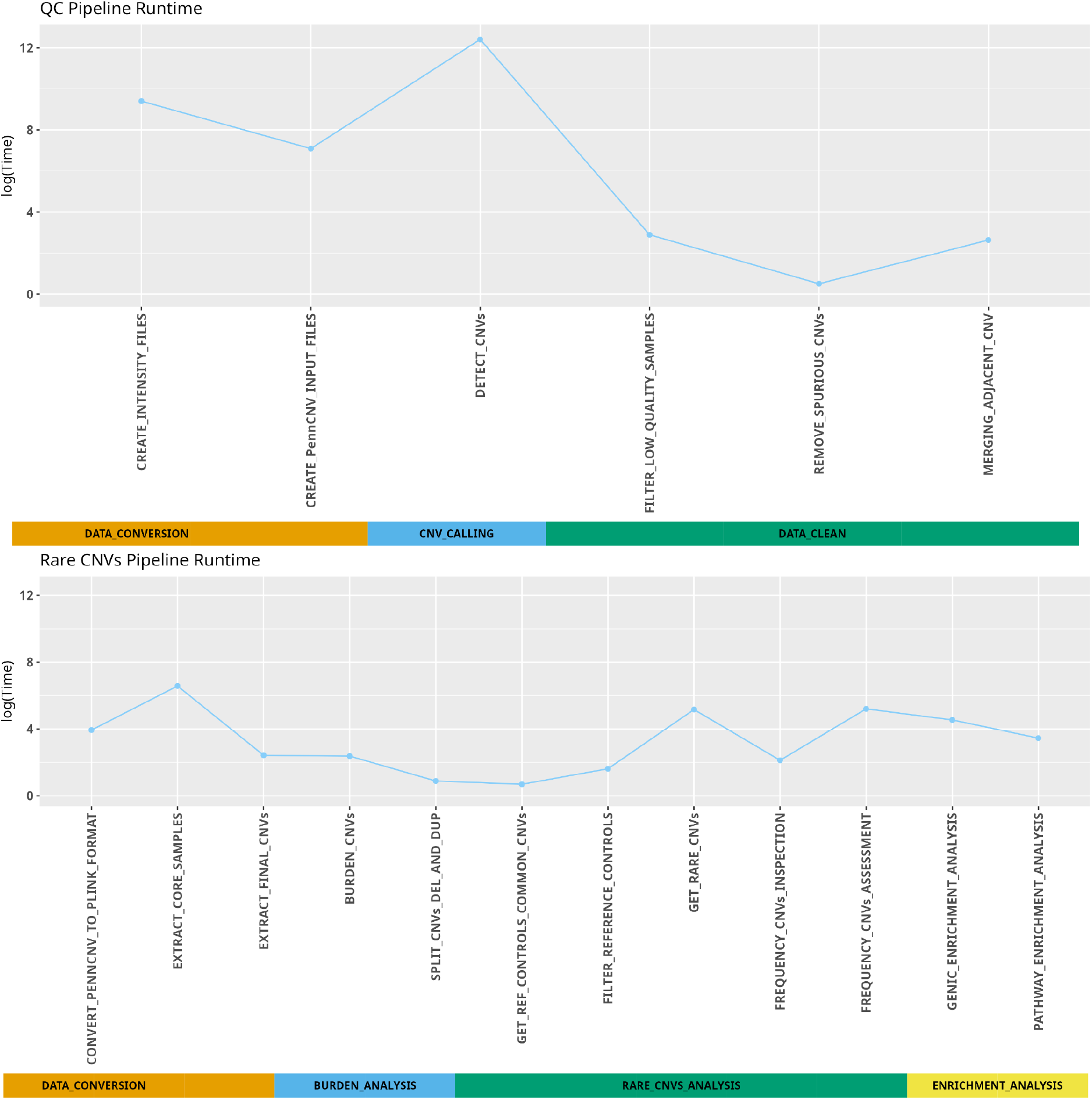
Pipeline performance. QC and rare-CNVs analysis for 6,112 samples, 700,079 markers (genotyping data from Illumina GSA), and 98,702 calls detected. Time in seconds, in logarithmic scale, is plotted for each module-rule. QC analysis runtime was 72.31 hours (260320 seconds), and rare-CNVs analysis runtime was 21.5 minutes (1290 seconds).

In the final stage of the CNV analysis pipeline, the Plink geneset-enrichment method is employed. This test compares the rate of CNVs impacting specific gene sets in cases versus controls, considering extreme biases in gene size and differences in CNV rate and size[26]. The pipeline includes two tests by default: the enrichment of genic CNVs (asking the question whether there is a general enrichment of genes among case CNVs), and the enrichment of pathway (or a predefined list of) genes, relative to all CNVs (determining whether there are a subset of genes enriched, relative to the whole genome). Both tests are based on a permutation test with N = 10,000 null permutations to generate empirical p-values (N can be modified inside the enrichment analysis module). The genomic coordinates of the genes, as well as the pathways to be tested are provided as configuration files to the pipeline. The enrichment test performs a generalized linear model-based (GLM-based) CNV burden test, and evaluates gene counts (GCNT), number of segments or CNVs (NSEG), and average size of CNVs (AVGKB) using logistic regression.

### External code and logs

A rule in a specific module can include inline code in Python or shell commands. However, extensive code within a single rule might hinder the module-rule modification. An external file (function.sh) containing shell functions used by some modules (Figure 2) is included with the pipeline utilities, making the inclusion or modification of external shell code clearer and simpler.

Both the QC pipeline and rare CNVs pipeline automatically generate the log text files (inside the logs directory) with relevant information for each module, such as number of samples included/excluded, number of calls filtered, burden summary and enrichment summary. Logs can be used to create a report including overall information as presented in Table 1 and supplementary Table 2 and 3.

**Table 1.**
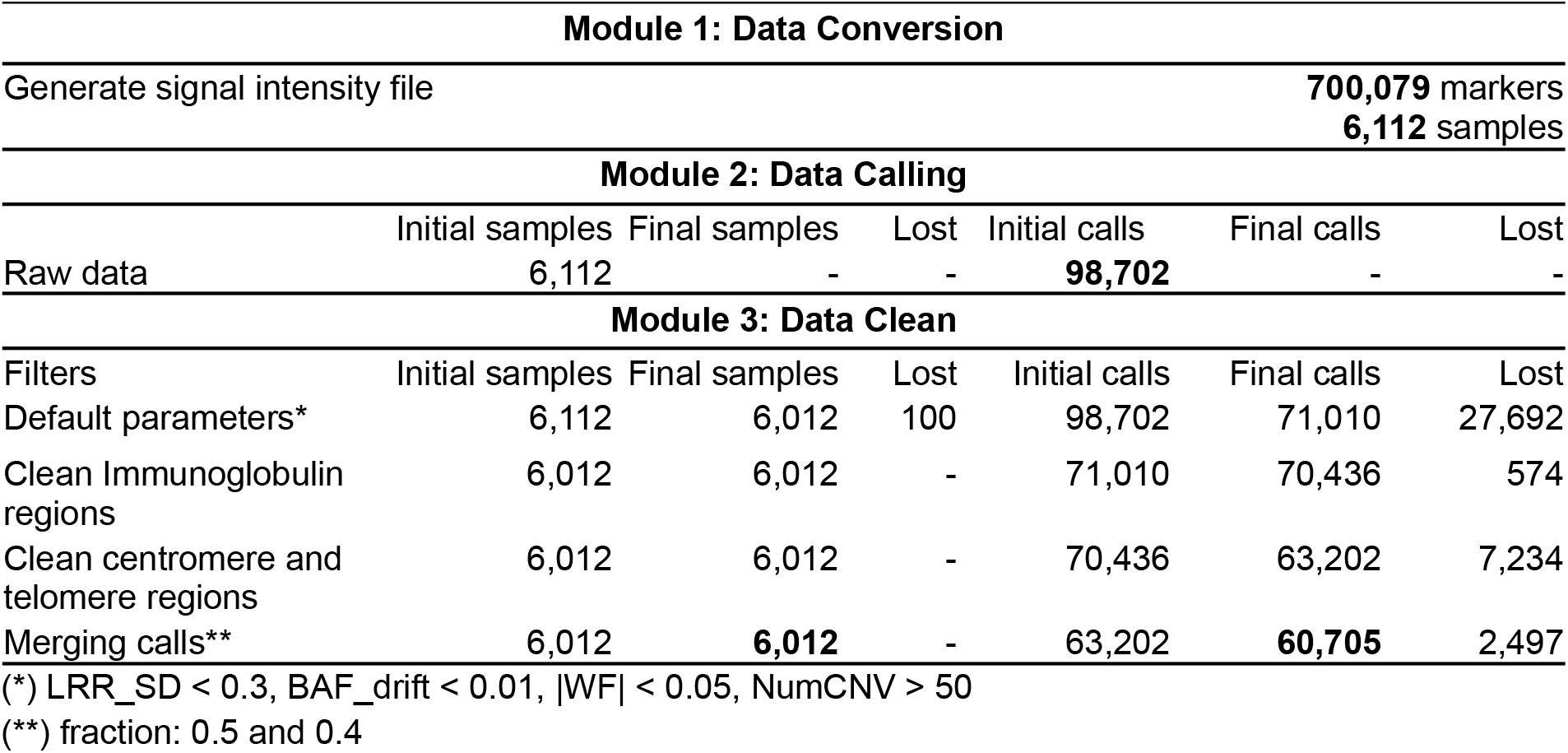
Calls and samples included and excluded at each module in QC pipeline. Final samples and calls, after QC, are in bold.

| Module 1: Data Conversion |  |  |  |  |  |  |
| --- | --- | --- | --- | --- | --- | --- |
| Generate signal intensity file |  |  |  |  | **700,079** markers | **6,112** samples |
| Module 2: Data Calling |  |  |  |  |  |  |
|  | Initial samples | Final samples | Lost | Initial calls | Final calls | Lost |
| Raw data | 6,112 | - | - | **98,702** | - | - |
| Module 3: Data Clean |  |  |  |  |  |  |
| Filters | Initial samples | Final samples | Lost | Initial calls | Final calls | Lost |
| Default parameters* | 6,112 | 6,012 | 100 | 98,702 | 71,010 | 27,692 |
| Clean Immunoglobulin regions | 6,012 | 6,012 | - | 71,010 | 70,436 | 574 |
| Clean centromere and telomere regions | 6,012 | 6,012 | - | 70,436 | 63,202 | 7,234 |
| Merging calls** | 6,012 | **6,012** | - | 63,202 | **60,705** | 2,497 |

(\*) LRR\_SD $< 0.3$ , BAF\_drift $< 0.01$ , |WF| $< 0.05$ , NumCNV $> 50$
(\*\*) fraction: 0.5 and 0.4

### Performance

This pipeline executes non-parallel tasks, although Snakemake can automatically determine which parts of the workflow can be run in parallel, decreasing the execution time of some modules. Figure 3 shows the runtime for both the QC pipeline and rare-CNV pipeline, for genotyping data (from Illumina GSA) of 6,112 samples, 700,079 markers, and 98,702 calls detected. The QC pipeline execution time, approximately 72 hours, takes most of the total time of the execution, especially modules which perform the data conversion from the signal intensity values to PennCNV, and CNV calling. It should be mentioned that these modules will be executed only on the first run. The downstream rules, directly involved with samples and calls quality, can be modified and the QC pipeline can be executed again, skipping the run of the previous modules which decreases the execution time substantially. Similar approach is applied for the rare-CNVs pipeline.

Due to the security requirements for personally identifiable data used in this performance testing, we used the TRE provided by the HUNT cloud secure solutions for scientific cloud computing (ntnu.edu/mh/huntcloud):

Operative system: Ubuntu 18.04.6 LTS (GNU/Linux 4.15.0-210-generic x86_64)

Architecture: x86_64

CPU op-mode(s): 32-bit, 64-bit

CPU(s): 32

Model name: Intel Core Processor (Broadwell, no TSX, IBRS)

CPU MHz: 2095.074

Memory: 64 GB

Total runtime: 72.67 hours.

Moreover, this pipeline can be run on a standard desktop computer. A basic test was performed using a small demo data (12 samples, 654,028 markers and 472 calls detected) downloaded from Illumina in an Ubuntu virtual machine (see test in RareCNVsAnalysis Github project):

Operative system: Ubuntu 22.04.4 LTS

Architecture: x86_64

CPU op-mode(s): 32-bit, 64-bit

Model name: 11th Gen Intel(R) Core(TM) i7-1165G7 @ 2.80GHz

Memory: 4 GB

Total runtime: 6.21 minutes

## Results and Discussion

We have generated a versatile pipeline for CNV detection and analysis from SNP arrays. To demonstrate the use of the pipeline we applied it to a case-control study in Addison’s disease where the results are presented in more details[27]. Samples were genotyped with Illumina Infinium Global Screening Array 1.0. CNVs were called and quality controlled using the pipeline. The box plots displaying PennCNV statistics values (LRR_mean, LRR_SD, BAF_mean, BAF_SD, BAF_drift and |WF|) were generated to assess the quality of the samples (Supplementary Figure 8). This plot illustrates samples meeting or failing the exclusion criteria based on the PennCNV QC threshold. Reduced overlap in the side-by-side box plots signifies a robust quality predictor (LRR_SD in this study). Furthemore, an abnormally high count of CNVs in a sample (NumCNV) suggests a low quality at a sample-level; such samples should be therefore excluded. The NumCNV threshold (> 50 in this study) can be established by inspecting the correspondence among samples failing or passing QC and the NumCNV (Supplementary Figure 9). After sample QC, potentially artificial CNV calls were removed from repeat-rich genomic regions such as HLA, telomeric, and centromeric regions, and then CNVs were merged to produce a set of high quality CNV calls (Supplementary Figure 10). Table 1 illustrates the main steps of the QC pipeline and the number of samples and CNVs included and excluded in each step.

After filtering samples and CNVs, the analysis of rare CNVs was executed. First, the PennCNV sample files were converted to Plink format and then, only unrelated 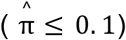 individuals of European descent were retained, as detailed in CNVs above 50 kb in length and spanning more than five markers were selected (default values can be changed in the pipeline parameter file) and a burden test for all CNVs was performed, which showed no significant differences in cases compared to controls in the four metrics, RATE (Number of segments), PROP (Proportion of samples with one or more segment), TOTKB (Total kb length spanned) and AVGKB (Average segment size) (Supplementary Figure 11). Continuing with the burden analysis, CNVs were classified into deletions and duplications, binned by length (by default 50 kb, 100 kb, 200, 500 kb and > 1,000 kb) and further the ratios in controls and cases were calculated (Supplementary Table 1). Once the burden analysis was finalized, the pipeline proceeded to rare CNV analysis, in which the rare deletions and duplications were extracted and evaluated for differences in frequency between cases and controls. For this study in particular, a subset of controls (200 individuals) previously selected were used as a reference to identify the common variants. Variants with count ≥ 4 (i.e. ≥ 2% carrier frequency) were classified as common variants. Subsequently, any CNVs overlapping at least at 50% of length with these common variants were excluded, to retain the rare variants with a frequency approximately < 2% (allele frequency < 1%). The frequency plot distribution for rare deletions and duplications, generated by the pipeline, enabled us to inspect these frequencies. The obtained frequencies fell within the predefined threshold for this study (Supplementary Figure 4 and 5). Next, the pipeline evaluated the cumulative distribution of CNV frequencies across five interval sizes (50-100, 100-200, 200-500, 500-1,000 kb and > 1,000 kb), estimating a two proportion test statistic and odds ratios (ORs). The results were then compiled in a summary file, alongside the forest plots (Table 2 and Table 3, and Supplementary Figure 6 and 7). The analysis which is described in detail in Artaza *et al*.*[27]* uncovered a higher frequency for the largest rare deletions (> 1,000 kb) among cases (n = 13/1182) compared to controls (n = 10/3810) (OR = 4.23, 95% CI 1.85-9.66, p = 0.0002). Finally, the pipeline performed the case-control gene-set enrichment test for two candidate gene-set list, primary immunodeficiency and congenital adrenal hypoplasia panels from the Genomics England PanelApp[28]. Based on the test results, no evidence supporting an overall enrichment of rare CNVs overlapping with immune related genes was observed[27] (Supplementary Figure 12).

**Table 2.** Overall rare deletions and duplications frequency distributions. This table shows data directly extracted from a summary text file. The table format can be adjusted by the user.

| CNV | Cases | Controls | Cases_freq | Controls_freq | P.value | OR | X95.CI | P |
| --- | --- | --- | --- | --- | --- | --- | --- | --- |
| DELs | 827 | 2615 | 0.6997 | 0.6864 | 0.4077 | 1.0646 | 0.9236,1.2269 | 0.3876 |
| DUPs | 721 | 2367 | 0.6100 | 0.6213 | 0.5073 | 0.9535 | 0.8339,1.0901 | 0.4857 |

**CNV:** CNV type, **Cases/Controls:** number of CNVs (deletions or duplications) in each cohort. **Cases\_freq/Controls\_freq:** CNVs frequencies, **P.value:** two proportion test p-value, **OR:** odds ratio, **X95.C:** confidence interval at 95%, **P:** odds ratio p-value associate.

**Table 3.** Rare CNV frequency distribution binning by size in cases vs. controls. This table shows data directly extracted from a summary text file. The table format can be adjusted by the user.

| Deletions |  |  |  |  |  |  |  |  |
| --- | --- | --- | --- | --- | --- | --- | --- | --- |
| Length | Cases | Controls | Cases_freq | Controls_freq | P.value | OR | X95.CI | P |
| 50KB_100KB | 435 | 1298 | 0.3680 | 0.3407 | 0.0911 | 1.1270 | 0.9837,1.2909 | 0.0846 |
| 100KB_200KB | 260 | 919 | 0.2200 | 0.2412 | 0.1435 | 0.8871 | 0.7586,1.0372 | 0.1331 |
| 200KB_500KB | 102 | 323 | 0.0863 | 0.0848 | 0.9174 | 1.0196 | 0.8078,1.2869 | 0.8703 |
| 500KB_1000KB | 17 | 65 | 0.0144 | 0.0171 | 0.6158 | 0.8407 | 0.4909,1.4397 | 0.5269 |
| 1000KB_1000000KB | 13 | 10 | 0.0110 | 0.0026 | 0.0005 | 4.2258 | 1.8481,9.6622 | 0.0002 |
| Duplications |  |  |  |  |  |  |  |  |
| Length | Cases | Controls | Cases_freq | Controls_freq | P.value | OR | X95.CI | P |
| 50KB_100KB | 297 | 1050 | 0.2513 | 0.2756 | 0.1078 | 0.8821 | 0.7597,1.0242 | 0.0998 |
| 100KB_200KB | 204 | 614 | 0.1726 | 0.1612 | 0.3773 | 1.0857 | 0.9125,1.2918 | 0.3536 |
| 200KB_500KB | 157 | 488 | 0.1328 | 0.1281 | 0.7077 | 1.0427 | 0.8596,1.2646 | 0.6712 |
| 500KB_1000KB | 48 | 150 | 0.0406 | 0.0394 | 0.9161 | 1.0328 | 0.7411,1.4391 | 0.8488 |
| 1000KB_1000000KB | 15 | 65 | 0.0127 | 0.0171 | 0.3614 | 0.7406 | 0.4207,1.3033 | 0.2960 |

**Length:** CNV length interval, **Cases/Controls:** number of CNVs (deletions or duplications) in each cohort. **Cases\_freq/Controls\_freq:** CNVs frequencies, **P.value:** two proportion test p-value, **OR:** odds ratio, **X95.C:** confidence interval at 95%, **P:** odds ratio p-value associate.

## Conclusion

We present an automated, flexible, and scalable bioinformatic pipeline tailored for rare CNV analysis in case-control studies. Array technology has undergone a tremendous growth in both quantity and content over recent years. Although genotyping data facilitate CNV analysis, the major challenges in the CNV analysis involve the management of large volumes of data, advanced bioinformatics, and complex data interpretation. Addressing this, a pipeline that streamlines analyses, systematizing tasks, while maintaining flexibility is indispensable. Our pipeline provides the fundamental steps for rare CNVs analysis, enabling automation of analyses while maintaining flexibility. Beyond the analysis of rare CNVs, the design principles using standardized modules render the pipeline reusable across a broad spectrum of bioinformatic analyses.

## Availability and requirements

Project name: Rare CNVs Analysis Pipeline. Project home page: https://github.com/haydeeartaza/RareCNVsAnalysis. Operating system(s): Linux, MacOS. Programming language: R, Shell Scripting, Python. License: MIT license. Any restrictions to use by non-academics: none. We used our in-house data to test the pipeline due to paucity of suitable publically available datasets.

## Supporting information

https://drive.google.com/file/d/1YvMJiP4EDZiUjbCukgRQWik822JuXeO8/view?usp=sharing

## Abbreviations

CNV: Copy number variation
SNP: Single nucleotide polymorphism
GWAS: Genome wide association study
LRR: Log R Ratio
BAF: B allele frequency
WF: Waviness Factor
NumCNVs: Number of called CNVs
PFB: Population frequency of B allele
PCA: Principal component analysis
MDS: Multidimensional scaling
IBD: Identity by descent

## Acknowledgements

Test was performed using digital laboratories in HUNT Cloud at the Norwegian University of Science and Technology, Trondheim, Norway. We are grateful for outstanding support from the HUNT Cloud community.

## Author Contributions

HA wrote the code, implemented the pipeline, performed the analysis, interpreted the results and wrote the manuscript. KL contributed with the code, gave software feedback and contributed to the manuscript. ASBW and ECR conceptualised and contributed to the manuscript. SJ and MV conceptualised, participated in supervision of the project, and wrote the manuscript. All authors have read and approved the final version of the manuscript.

