## Supplementary material for "Rare Copy Number Variant analysis in case-control studies using SNP Array Data: a scalable and automated data analysis pipeline": https://drive.google.com/file/d/1YvMJiP4EDZiUjbCukgRQWik822JuXeO8/view?usp=sharing

**Supplementary Figures and Tables**

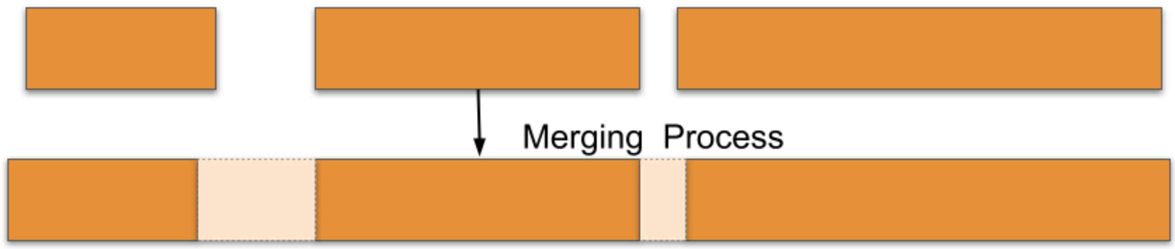

Supplementary Figure 1. Merging CNV calls. PennCNV has a tendency to divide large CNVs into smaller fragments. The pipeline combines neighboring CNVs into segments, with the default setting merging adjacent calls when the fraction of the gap length between two calls compared to the total length of the resulting merged call is <20%.

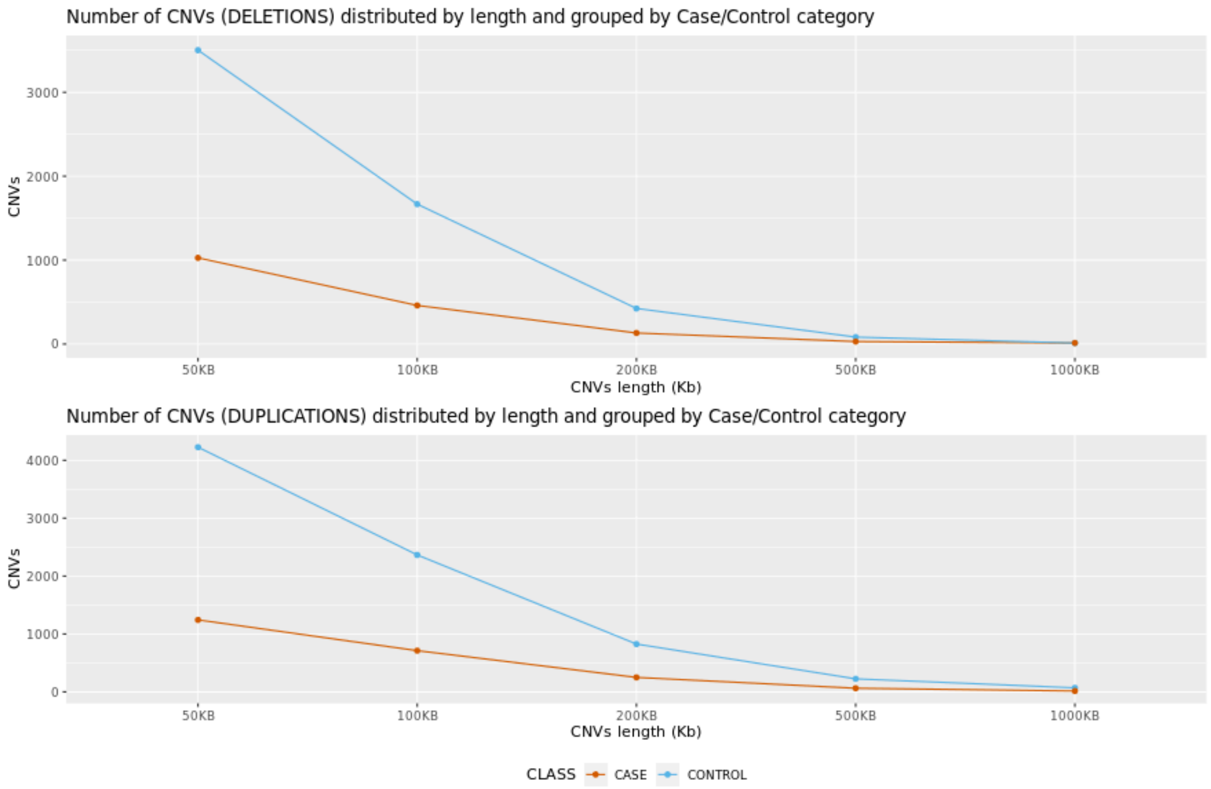

Supplementary Figure 2. Cumulative number of CNVs by size. A good sanity check early on in the study can prevent inaccurate results and save time spent running the analysis. This plot confirms the natural accumulation tendency of CNVs according to their length and the uniform CNVs distribution in cases and controls.

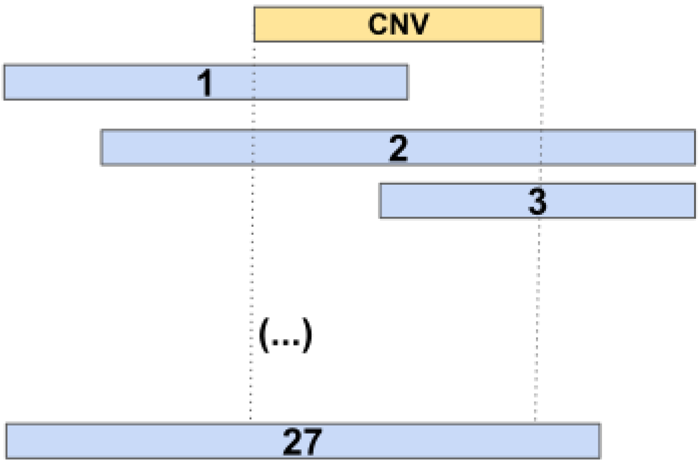

Supplementary Figure 3. CNV frequency calculation. For each CNV, the pipeline calculates the total number of CNVs that overlap with this CNV, including itself. The frequency is subsequently determined based on these counts. A 50% union overlap criterion is employed to ensure symmetry (i.e. if A overlaps B, then B must overlap A). In this example, the count for the target CNV region (in yellow) will be 28, indicating that 28 individuals harbor approximately the same CNV (in yellow) The frequency for this CNV is then calculated to be 28 divided by the total number of samples.

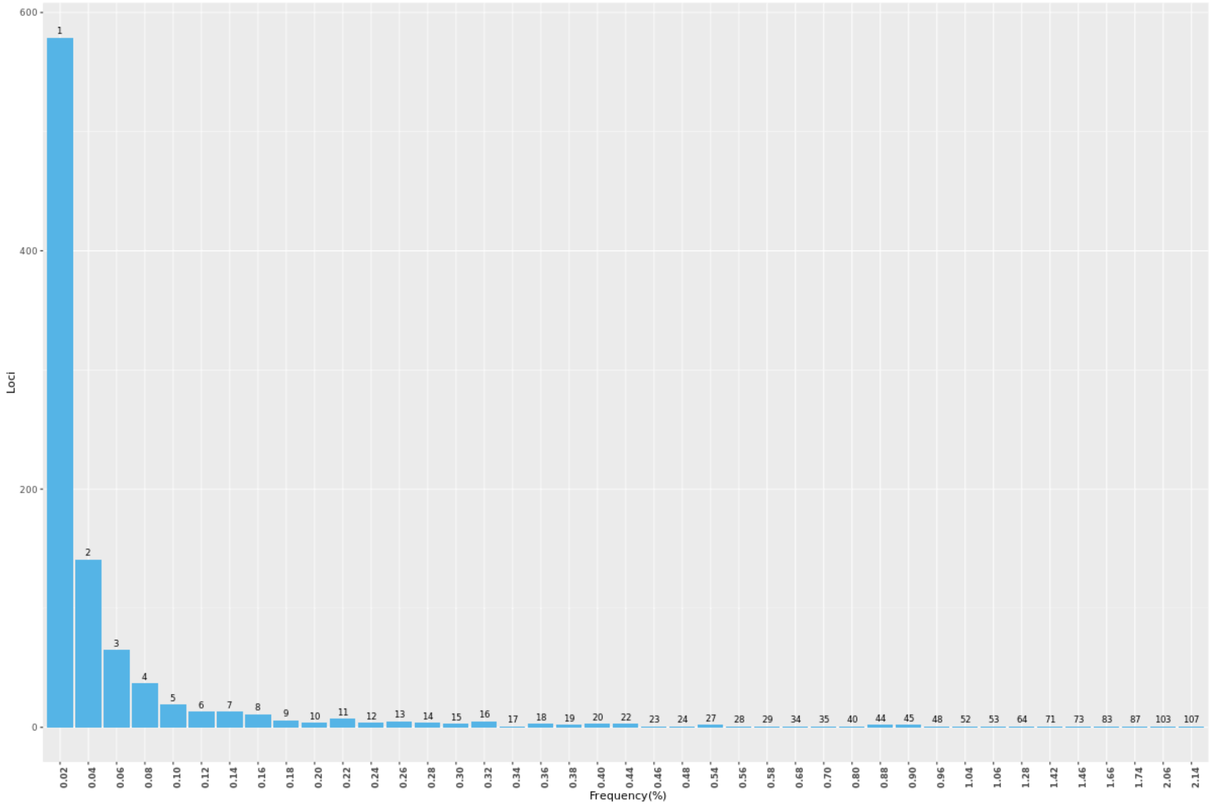

Supplementary Figure 4. Rare deletions frequency distribution. The Y axis shows the number of loci (regions), the X axis shows the frequency, and the number in the bar shows the amount of deletions in each locus that this frequency represents. E.g. 141 loci or regions (second bar size) contains 2 deletions (number in the bar) which represents ~0.04% of the frequency.

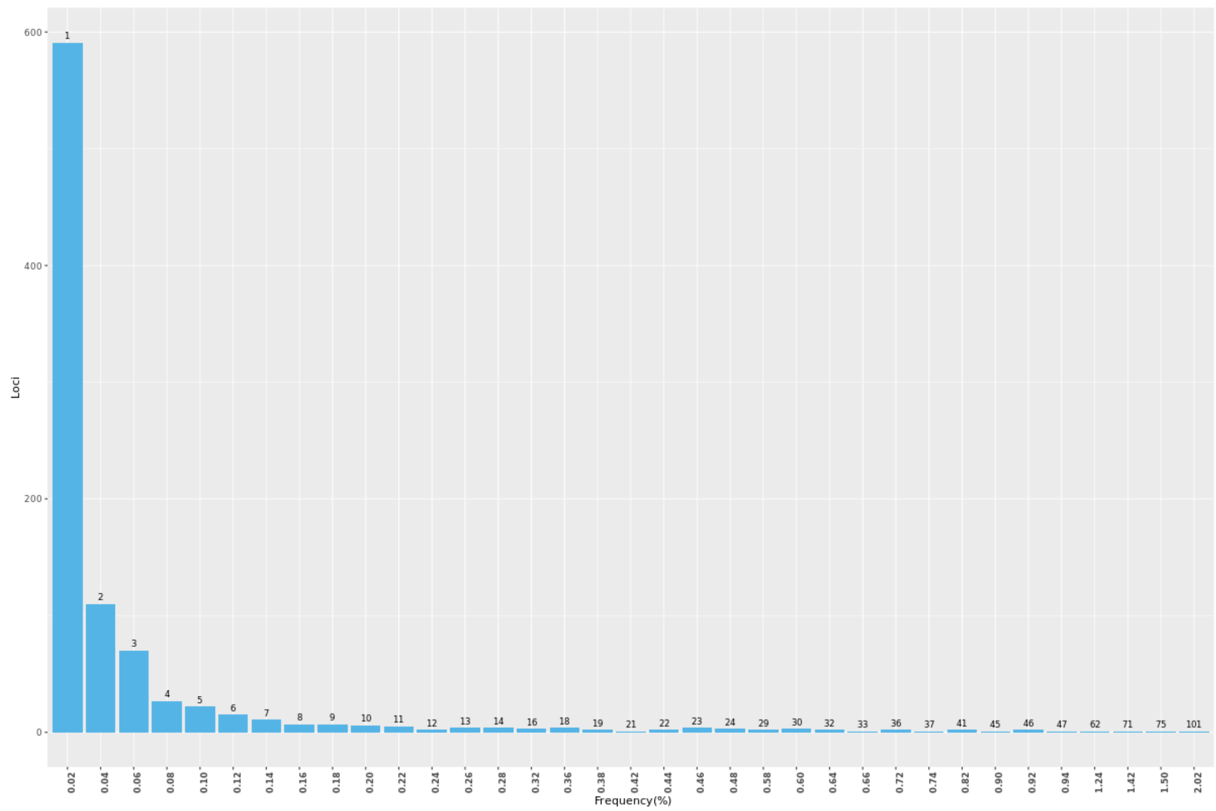

Supplementary Figure 5. Rare duplication frequency distribution. The Y axis shows the number of loci (regions), the X axis shows the frequency, and the number in the bar shows the amount of duplications in each locus that this frequency represents. E.g. ~600 loci or regions (first bar size) contains 1 duplication (number in the bar) which represents ~0.02% of the frequency.

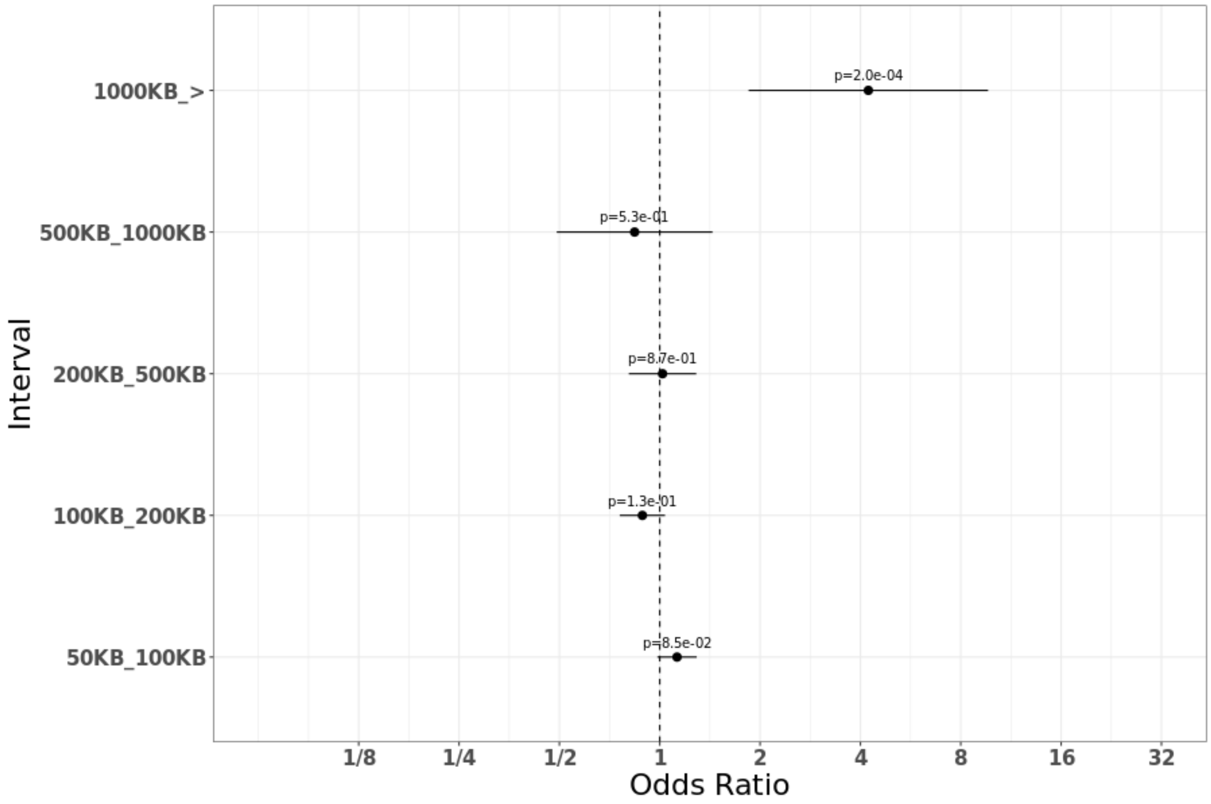

Supplementary Figure 6. Rare deletion frequencies distributed by size. Frequencies are shown by closed circles and whiskers represent the 95% confidence interval. The P-value associated is located above the line. For long deletions (> 1000 kb) a significant difference in the frequency was found between cases and controls.

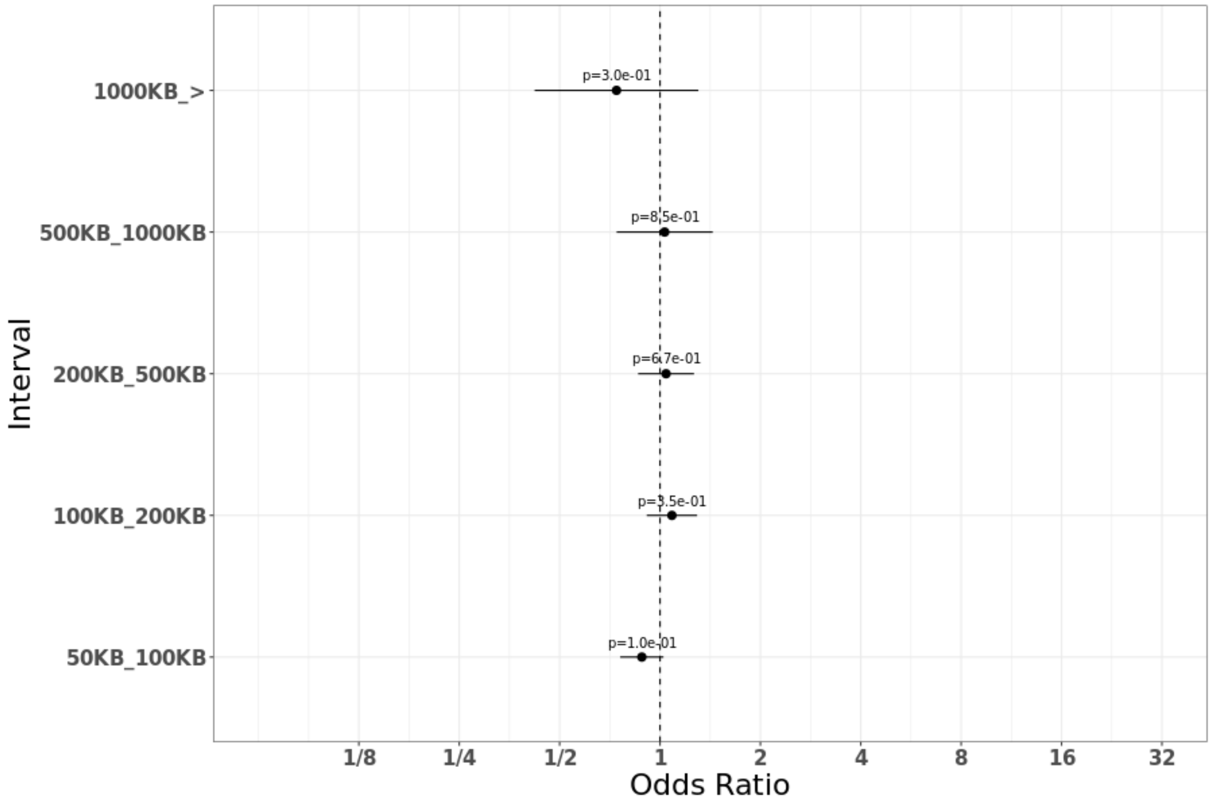

Supplementary Figure 7. Rare duplications frequencies distributed by size. Frequencies are shown by closed circles and whiskers representing the 95% confidence interval. The P-value associated is located above the line. No significant difference in the frequencies were detected.

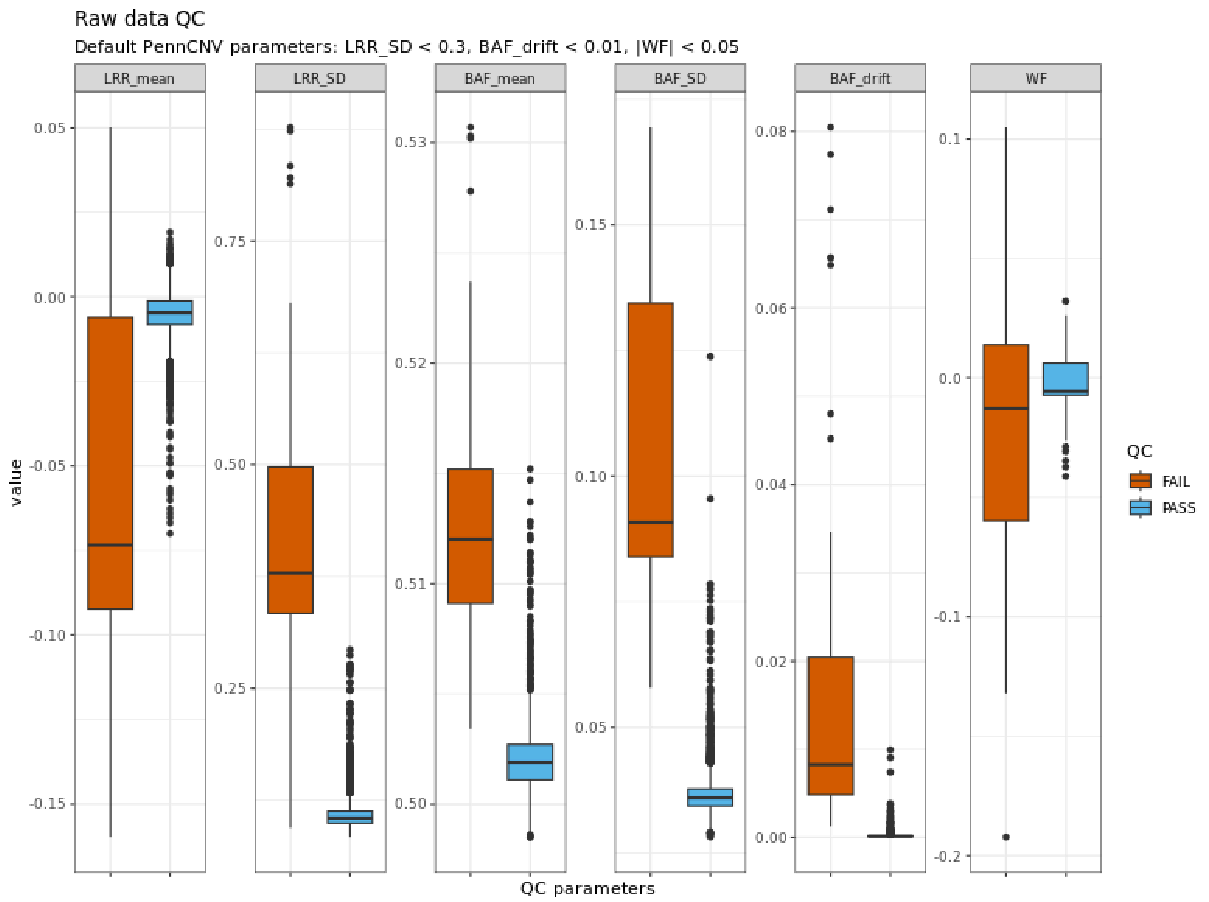

Supplementary Figure 8. Sample quality parameters. Red boxes show samples which fail the inclusion criteria based on the PennCNV QC threshold (LRR_SD < 0.3 & BAF_drift < 0.01 & |WF| < 0.05). Blue boxes show samples which pass the quality control.

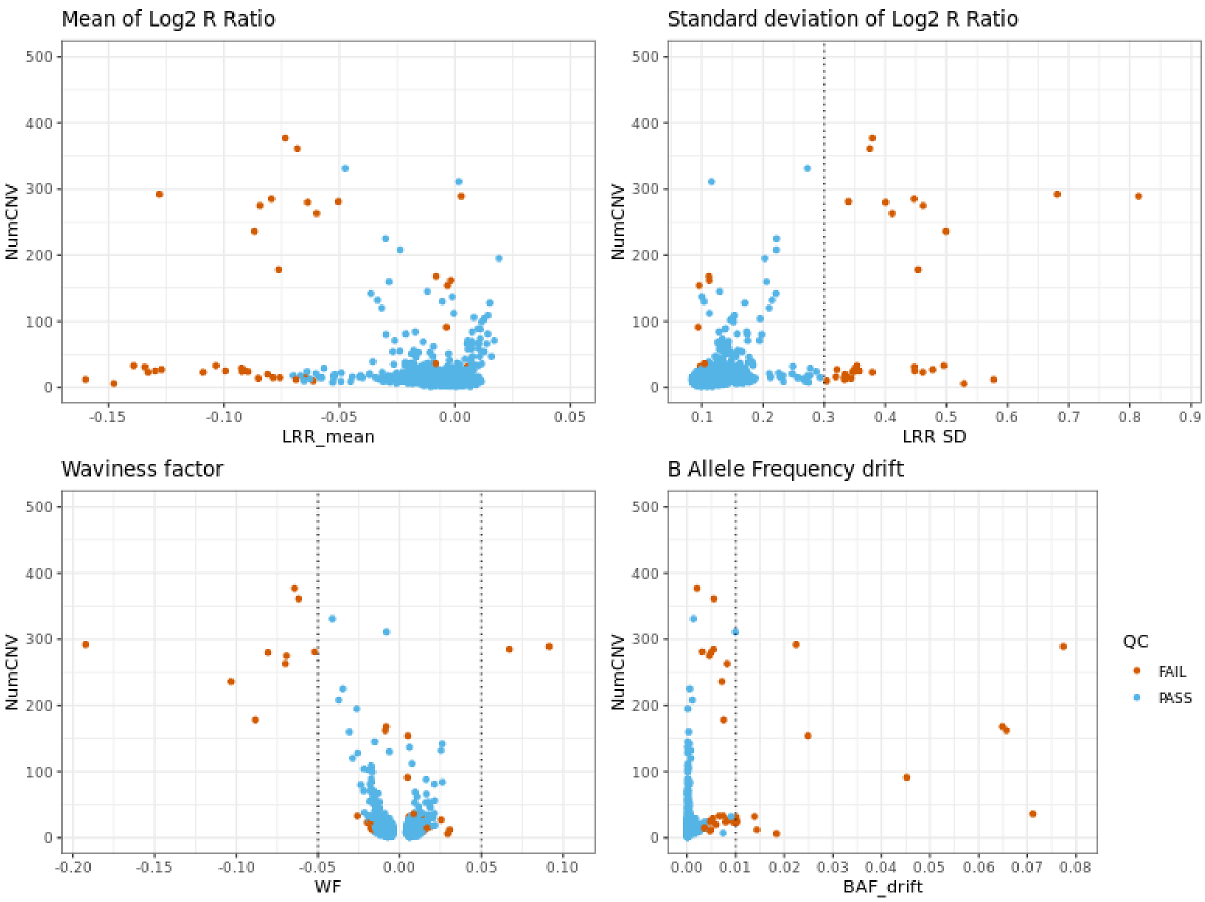

Supplementary Figure 9. The distribution of the number of CNVs per sample. Samples with an excessive number of CNVs should be considered for exclusion because it can indicate low data quality. A threshold for the number of CNVs per sample (NumCNV) can be defined through visual inspection, considering its distribution around the exclusion criteria threshold values based on PennCNV statistics. In the Addison's study (Artaza et al. doi:10.3389/fimmu.2024.1374499), samples with NumCNV >50 were removed.

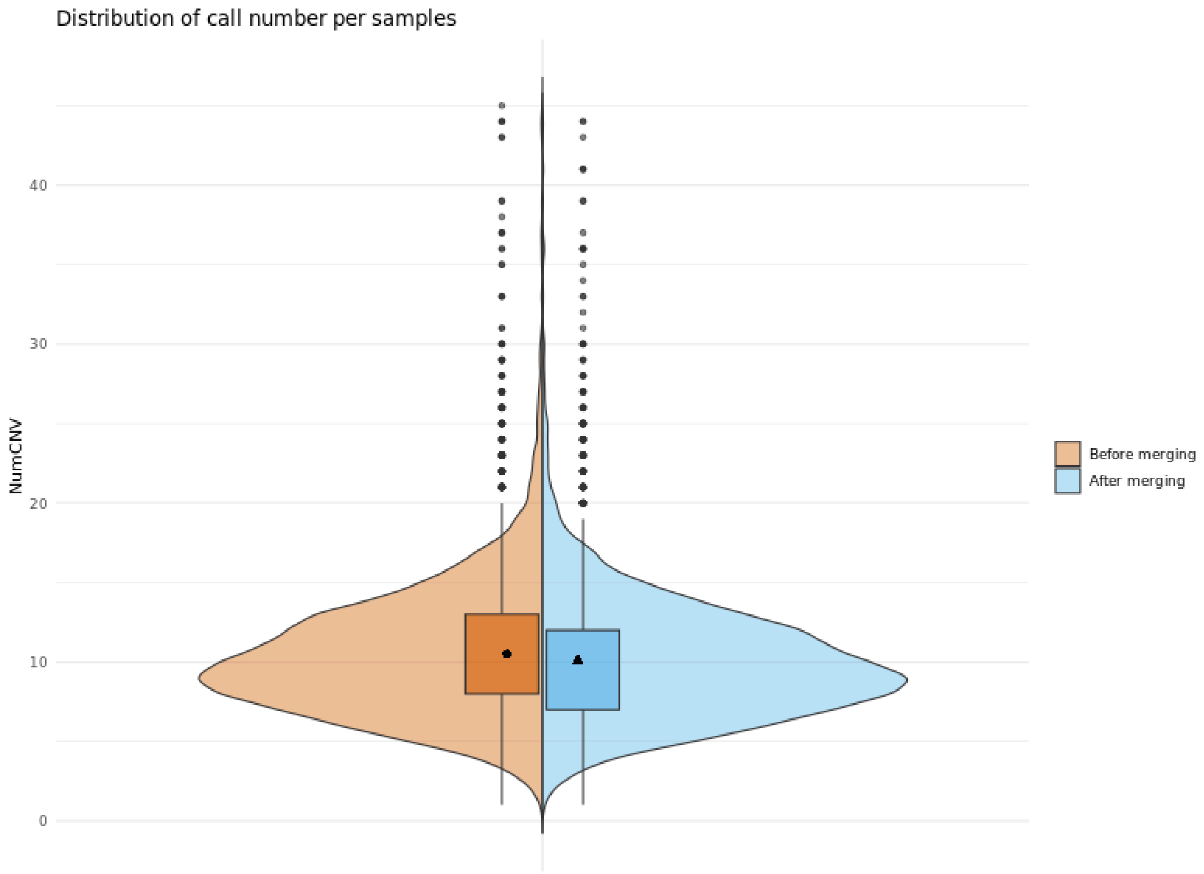

Supplementary Figure 10. The distribution of the number of calls per sample before and after the merging process. Following this process the number of calls per sample decreases as adjacent calls are merged into single calls.

(a) study_cnv.summary.mperm file

| CHR | SNP | EMP1 |
|---|---|---|
| S | RATE | 0.567443 |
| S | PROP | 0.715528 |
| S | KBTOT | 0.339566 |
| S | KBAVG | 0.284772 |
| S | GRATE | 1 |
| S | GPROP | 1 |
| S | GRICH | 1 |

(b) study_cnv.grp.summary file

| TEST | GRP | AFF | UNAFF |
|---|---|---|---|
| N | ALL | 2270 | 7728 |
| RATE | ALL | 1.92 | 1.927 |
| PROP | ALL | 0.8477 | 0.8536 |
| TOTKB | ALL | 362.6 | 356.8 |
| AVGKB | ALL | 160.5 | 157.2 |

Supplementary Figure 11. Basic burden analysis outcome**.** The figure shows two files generated in the basic burden analysis for the four metrics RATE (Number of segments), PROP (Proportion of sample with one or more segment), TOTKB (Total kb length spanned), AVGKB (Average segment size): (a) permuted results for segment test, includes the standard empirical p-value; and (b) group cases (affected, AFF) and controls (unaffected, UNAFF) summary statistics.

| TEST | BETA | P |
|---|---|---|
| GCNT | 0.0146582 | 0.400695 |
| NSEG | -0.0204375 | 0.786693 |
| AVGKB | 0.000311012 | 0.320859 |

Supplementary Figure 12. Enrichment test analysis outcome. The enrichment test consists of a GLM-based CNV burden analysis, yielding logistic regression parameters associated with gene counts (GCNT), number of segments or CNVs (NSEG), the average size of CNVs (AVGKB), and their respective P-values.

Supplementary Table 1. Burden analysis.

| **Deletions** | | | |
|---|---|---|---|
| | length | num_CNVs | ratio |
| Controls | 50KB | 3502 | 0.87 |
| | 100KB | 1667 | 0.42 |
| | 200KB | 424 | 0.11 |
| | 500KB | 83 | 0.02 |
| | 1000KB | 12 | 0.00 |
| Cases | 50KB | 1027 | 0.87 |
| | 100KB | 460 | 0.39 |
| | 200KB | 132 | 0.11 |
| | 500KB | 30 | 0.03 |
| | 1000KB | 13 | 0.01 |
| **Duplications** | | | |
| | length | num_CNVs | ratio |
| Controls | 50KB | 4226 | 1.05 |
| | 100KB | 2365 | 0.59 |
| | 200KB | 826 | 0.21 |
| | 500KB | 225 | 0.06 |
| | 1000KB | 70 | 0.02 |
| Cases | 50KB | 1243 | 1.05 |
| | 100KB | 712 | 0.60 |
| | 200KB | 250 | 0.21 |
| | 500KB | 63 | 0.05 |
| | 1000KB | 15 | 0.01 |

CNVs (deletions and duplications), distributed by length in controls and cases. [**num_CNVs]**: total number of CNVs in each interval length, [**ratio]**: number of CNVs divided by number of cases or controls respectively.

Supplementary Table 2. Sample and call numbers in the rare CNV pipeline.

| | Total samples | Cases | Controls | Males | Females | CNV calls |
|---|---|---|---|---|---|---|
| Raw data | 6112* | 1526 | 4471 | 3155 | 2946 | 98702 |
| High quality **core** samples | 5192 | 1182 | 4010 | 2705 | 2487 | 52144 |
| Samples with CNVs > 50KB & 5 SNPs | **4425** | 1002 | 3423 | 2302 | 2123 | **9998** |

Includes information on the cohort, gender and CNV calls. These details are documented in the log file of the data conversion module.

(*) 7 samples without classification: case/control

Supplementary Table 3. Rare CNVs extraction strategy.

| Type | Step | | CNVs | Samples |
|---|---|---|---|---|
| All | 1. | Initial core data (high quality samples and CNVs > 50KB and covered by at least 5 SNPs) | 9998 | 4425 |
| | 2. | Define 200 reference controls | 442 | 200 |
| | 3. | Remove 200 reference controls defined in (2) from initial data | **9556** | **4225** |
| DELETIONS | 4. | Initial deletions in core data (high quality samples and CNVs > 50KB covered by at least 5 SNPs) | 4351 | 2903 |
| | 5. | Common deletions (frequency >= 2%) from 200 reference controls | 46 | 50 |
| | 6. | Identify CNVs in (4) overlapping at least 50% with common variants (reciprocal) | 909 | 856 |
| | 7. | Remove common deletions from the dataset in (6) | **3442** | 2470 |
| DUPLICATIONS | 8. | Initial duplications in core data (high quality samples and CNVs > 50KB and covered by at least 5 SNPs) | 5205 | 3167 |
| | 9. | Common duplications (frequency >= 2%) from 200 reference controls | 119 | 99 |
| | 10. | Identify CNVs in (8) overlapping at least 50% with common variants (reciprocal) | 2117 | 1746 |
| | 11. | Remove common duplications from the dataset in (10) | **3088** | 2212 |

Following QC and filtering CNVs > 50 kb and covered by more than five probes, 200 reference control individuals (specific thresholds for this example) are extracted from the control cohort to identify common variants (frequency >= 2%). These individuals are then removed from the control group, along with their respective CNV calls. After that, rare deletions and duplications are extracted, with CNVs overlapping at least 50% of the reference common variants being excluded. These details are documented in the log file of the rare CNV analysis module.
